# Build me up to break me down: Frothed spawn in the sandpaper frog, *Lechriodus fletcheri*, is formed by female parents and later broken down by their offspring

**DOI:** 10.1101/2020.02.06.937409

**Authors:** John Gould

**Affiliations:** School of Environmental and Life Sciences, University of Newcastle, Callaghan, 2308 NSW, Australia

## Abstract

Several genera of anuran amphibians deposit their eggs within mucous secretions that have been aerated by the parents to produce a foam or bubble spawn body. This is a dynamic medium for embryo development given that is gradually breaks down over time, and one that has been hypothesised to serve a variety of purposes including protecting embryos from external stresses. In this study, I provide additional details of bubble spawn production in the sandpaper frog, *Lechriodus fletcheri*. Field and laboratory observations showed that females aerate spawn while in inguinal amplexus, using flanged fingers to transport air bubbles into the mucous. While the frothed spawn is initially resistant to breakup, it gradually loses bubbles and flattens out into a film. This temporal shift in structure is likely to be adaptive, as the resultant increase in surface area allows embryos to come in direct contact with the open water, which may accommodate their increased oxygen demands or facilitate hatching. I provide evidence that this process is controlled by the residing embryos, given that spawn in their absence does not breakdown, highlighting the ability of offspring to modify their immediate environment even before hatching to ensure conditions remain suitable for their changing needs.

## Introduction

A great diversity of ovipositional modes have evolved among amphibians (Duellman and Trueb 1986; Cockran and Thoms 1996; Stebbins 2003; Altig and McDiarmid 2007). This includes the release of eggs one or a few at a time to complete development alone or in loose egg aggregations, their deposition into more defined aggregations with their outer jelly layers in close contact, or their embedment within a matrix composed of mucous secretions or turgid sac. Each ovipositional mode will influence the manner in which offspring interact with their external environment and, as such, may play a critical role in determining the chances of their survival by minimising the effect of external stresses (Stamp 1980). Yet for many species, accurate and detailed descriptions of oviposiitoning behaviour and subsequent egg clutch structures remain to be reported in the literature.

Among several anuran genera, adults will deposit their eggs in mucous that has been aerated to produce either a foam or bubble spawn body, both of which are referred to as a froth ‘nest’ (Frost 1985). This behaviour has evolved independently among both *ranoid* (*Rhacophoridae*, *Hyperoliidae*) and *hyloid* frogs (*Myobatrachidae*, *Leptodactylidae*), with the manner of aeration and the resultant form of the frothed spawn varying immensely within and between groups. For example, spawn production may be aquatic or terrestrial, with frothing of precursor fluid facilitated by one or both parents, resulting in the production of spawn composed of either small or large bubbles. Mucous secretions may be physically whipped by movement of the adult’s legs or by pairs jumping onto the spawn to produce foam spawn (Heyer and Rand 1977; Bastos *et al.* 2010), or aerated by the transport of air bubbles by movement of the adult’s legs or via release through the nostrils of pairs to form bubble spawn (Tyler 1989; Hödl and Haddad 1997). Given the inclusion of air, the frothed spawn body will float on the surface of the water through some or all stages of embryo development. As such, it has been suggested to be a first evolutionary step away from the reliance on free-standing water and towards terrestrial egg deposition (Heyer 1969; Martin 1970). However, this behaviour may serve a variety of adaptive purposes, producing an incubation medium that serves to reduce oxygen competition among embryos (Seymour and Roberts 1991; Seymour and Bradford 1995; Hödl and Haddad 1997), or to protect embryos from thermal damage, desiccation and predation given the buffering properties of the inflated structure (Heyer 1969; Gorzula 1977; Dobkin and Gettinger 1985; Hödl 1986; Downie 1988).

For the Australian *Myobatrachid* anurans, bubble spawn production occurs across several genera including *Adelotus*, *Limnodynastes*, *Lechriodus*, *Heleioporus* and *Philoria* (Tyler and Davies 1979). In general, frothing is known to be performed by the female of aquatic species while in inguinal amplexus, her front legs churning the surrounding water to trap air bubbles underneath the gradually emerging eggs and associated mucous secretions that compose the spawn. Currently, the adaptive benefits of this behaviour for offspring survival remain poorly known in this group. Exacerbating the lack of investigation into this behaviour may be the lack of natural history data for many species, including the behaviours involved in amplexus and froth production, as well as the subsequent structure of the frothed spawn throughout embryo development. The present study describes aspects of oviposition mode in the sandpaper frog, *Lechriodus fletcheri*. This species exploits highly ephemeral pools for egg deposition, creating a bubble spawn body that deteriorates as the embryos develop (Anstis 2017). Field and laboratory observations in the current study provide insights into the behaviour of *L. fletcheri* parents during amplexus, their specific contributions to foam spawn production, and the structure of the frothed spawn body through till embryo hatching. Evidence is also presented which suggests that it is the embryos themselves that catalyse the gradual deterioration of the spawn body over the course of their development, as a likely means to provide direct access to open water prior to hatching.

## Methods

Field observations for this study occurred within the Watagan Mountain Range, New South Wales, Australia (33° 0’ 18.396” S, 151° 26’ 23.4312” E). Within this system, *L. fletcheri* has come to exploit ephemeral pools that form near dirt roads during periods of rainfall. Over four consecutive seasons (2014-15, 2015-16, 2016-17, 2017-18), approximately 80 pools were regularly surveyed at night for the presence of adults and spawn by scanning the water and surrounded leaf litter for their presence.

### Amplectant and ovipositing behaviour

Adults found in amplexus were photographed *in situ* before being collected and photographed to obtain detailed information on the positioning of pairs when coupled. Most adults were released back to their original point of capture within a few minutes. However, four amplectant pairs were collected for a companion study on one survey night (21 February 2018) and video-recorded in the laboratory to more closely investigate the process of spawn production. Pairs were collected in separate containers (170 × 120 × 70 mm) before being transferred to large tubs (350 × 210 × 210 mm) that contained a small amount of damp forest leaf litter. Pairs were supplied with a small container (170 × 120 × 70 mm) filled with aged tank water to allow for egg deposition. Containers were filled to a depth of approximately 50-60 mm so that pairs did not make contact with the bottom. Videos of 5 minute length were periodically taken of pairs from a dorsal view over the night of capture, making sure to prevent disturbing pairs while in amplexus. Videos were analysed to determine the sequence of events that occurred during egg release and froth production. Following amplexus, the front hands of adult individuals were photographed under 1000x magnification using a stereo-microscope mounted DAGE-MTI camera with Leica LAS EZ software V4.0.0 (Leica Microsystems) to examine hand structures used in amplexus (particularly nuptial excrescences or ‘pads’ in males) and spawn production (particularly flanged fingers in females) which are known to be present in this species (Tyler and Davies 1979).

### Spawn structure

Spawn found during field surveys were photographed and aged based on the developmental stage of the residing embryos (Gosner 1960). The mean circular diameter of fresh spawn within 24 hours of being deposited was measured from photographs in the program ImageJ (Schneider *et al.* 2012). Egg counts were also made from 64 spawn collected from the field across all breeding seasons. A small section of spawn material was also removed from three of the collected spawn and photographed under 1000x magnification using a stereo-microscope as before, with mean egg and bubble diameter measured from photographs in the program ImageJ.

In addition, two freshly laid spawn, including one which contained no eggs, were collected on one survey night (23 December 2015) and brought back to the laboratory to more closely observe changes in the structure of the frothed spawn over time. To prevent desiccation, these spawn were transported back to the laboratory in separate containers (170 × 120 × 70 mm) filled with a small amount of natal pond water. Spawn were kept in these containers upon arrival and allowed to develop under room temperature conditions away from direct sunlight.

All measurements have been made in terms of the mean and standard deviation (SD). All photographs and videos for this study besides microscopic images were taken using an iPhone 6 (Apple Corporation, Cupertino, California, United States).

## Results

### Amplexus and oviposition behaviour

A total of 58 amplecting pairs were recorded across the four breeding seasons. Each pair was found at the inner edge of the pond or floating at the surface of the water in inguinal amplexus, slightly tilted upwards in the water column so that only the head of the female and both the head and dorsal surfaces of the male were above water. Each male was coupled to a female on top, with both his hands clasped around the abdomen of the female immediately above her hind legs. The lateral side of both hands of the male were held firmly against the female’s abdomen, nearly completely wrapping around the female in a complete ‘hug’. Individuals were not aligned in each pair, with the cloaca and hind legs of the male seated below that of the female, and the snout of the male seated just below the eyes of the female. A small number of pairs (n ≈ 10) were found during spawn body construction. In these cases, the vast majority of the spawn trailed behind each pair, with some traces of white, frothed secretions surrounding their bodies and facial regions (Fig. 1).

**Figure 1.**
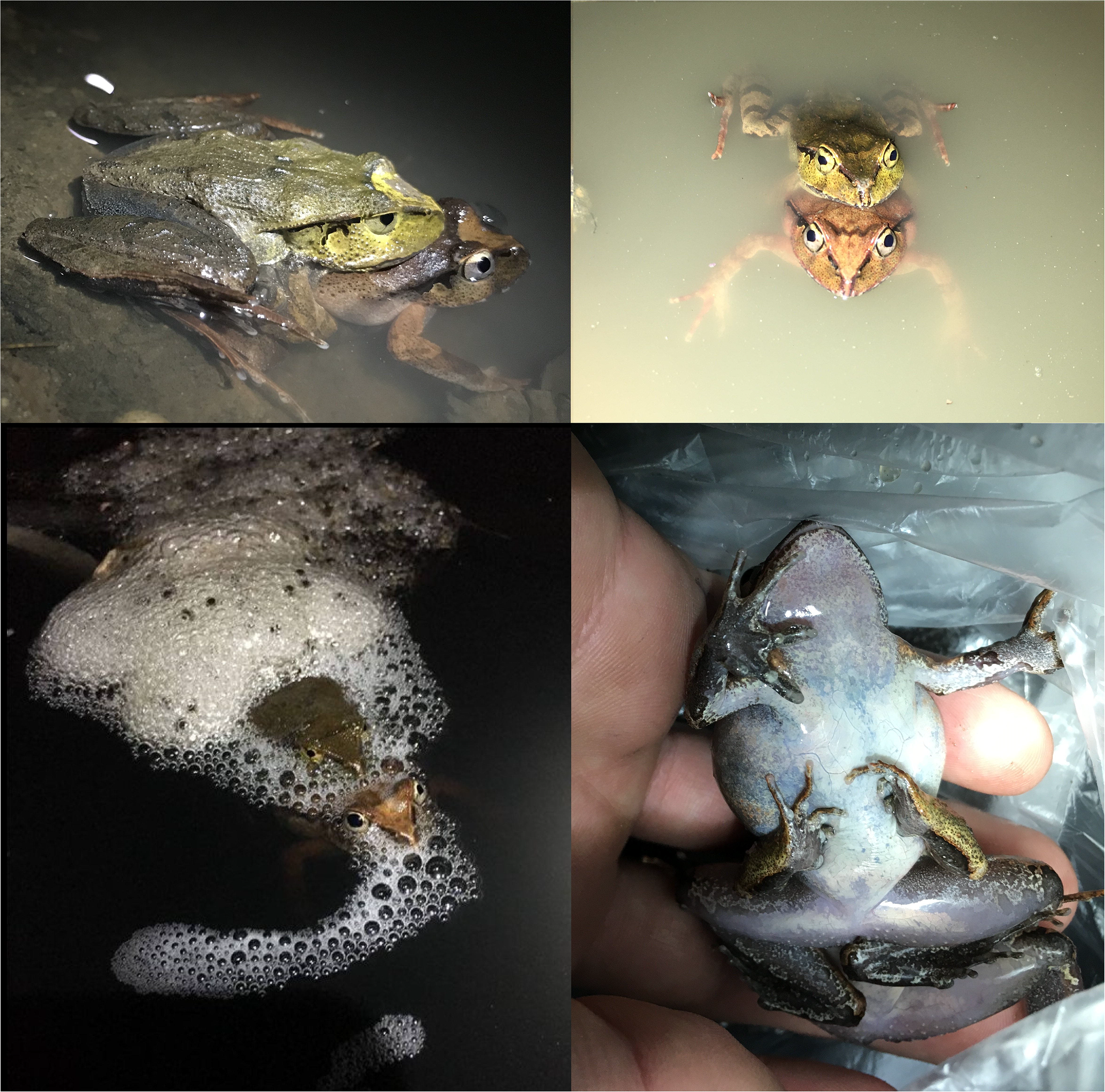
Amplecting behaviour of *Lechriodus fletcheri* adults at breeding pools within the Watagan Mountain Range, NSW, Australia. Photographs show the positioning of pairs from the side (top left), the front (top right) and from beneath (bottom right), as well as the positioning of pairs during spawn production (bottom left). Photographs are not to scale.

Of the four amplectant pairs transported back to the laboratory, video recordings of the ovipositing behaviour of one pair was obtained. During egg and mucous release, this pair remained immobile at the surface of the water, slightly tilted upwards in the water column in a manner similar to pairs found in the field. Only the female participated in beating mucous secretions into froth. She achieved this by initially paddling her front legs from her sides and near the surface of the water in a ‘windshield wiper’ motion, then inwards and downwards towards her abdomen, transporting air bubbles into the growing spawn body from underneath. Paddling motions were constant and regular, occurring every two seconds without rest, with the female’s head remaining above water and no involvement of the hind legs. Throughout this process, the male remained relatively motionless. Over a five minute interval, the spawn body behind the pair gradually increased in size, with older frothed material moving away from the pair as new material was released by the female. Traces of frothed mucous were present around the perimeter of the pair over this period. A video recording of this event can be found at https://archive.org/details/l.fletcheriamplexus (Supplementary Material).

The hands of adult *L. fletcheri* males were comprised of four fingers, along with a small, raised nub behind the inner-most finger. Nuptial pads were present on the lateral side of the inner two fingers of both hands, as well as on the raised nub. These pads were dark in pigment, composed of many small, conical projections, and extended from the base of each finger to just behind the toepad. The hands of adult females were also comprised of four fingers but lacked any apparent nub. Female fingers were relatively longer and thicker than those of males, with the outer two fingers of both hands flanged to form flattened, curved structures resembling scoops (Fig. 2).

**Figure 2.**
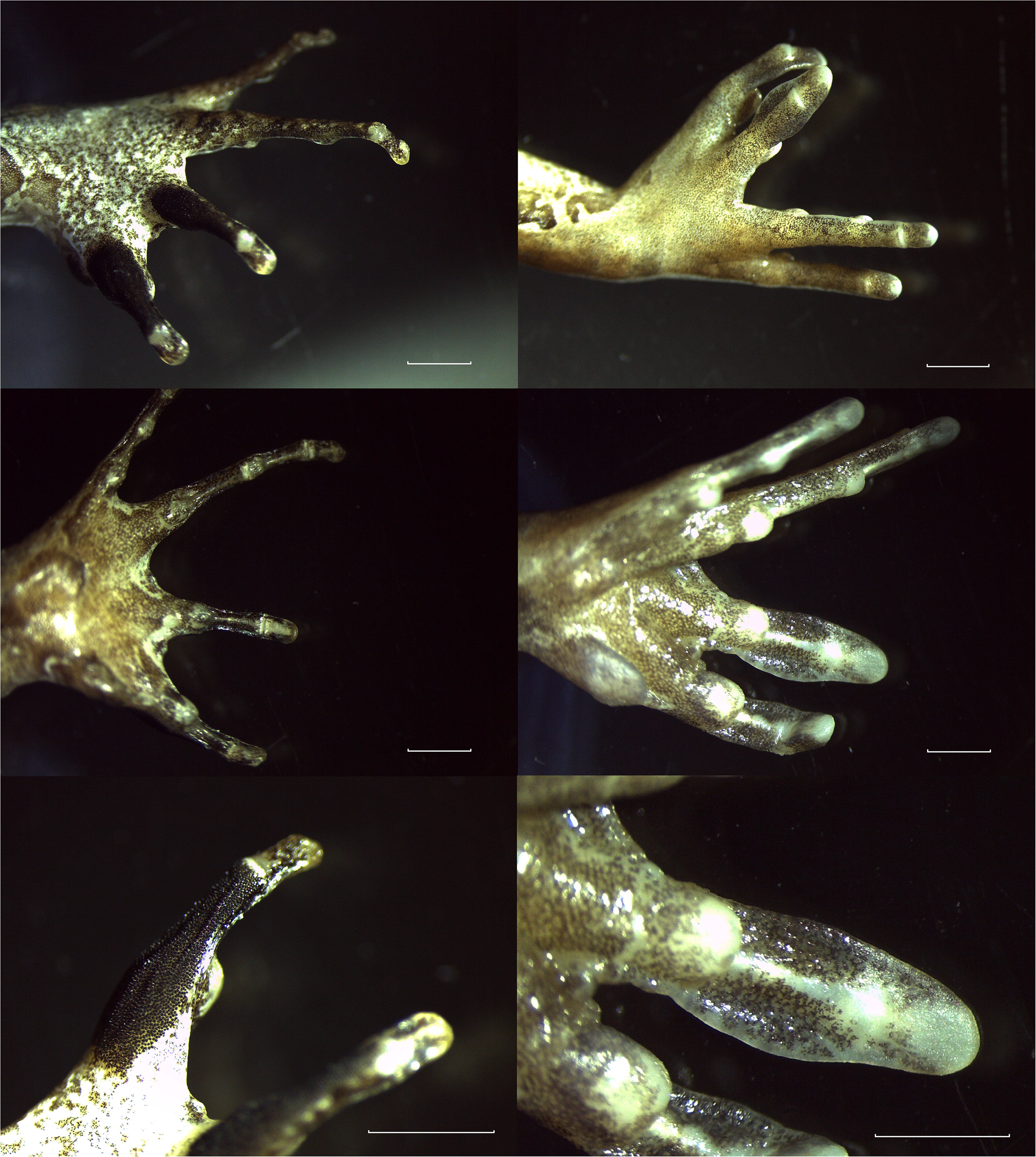
The morphology of reproductively active male and female *Lechriodus fletcheri* adults. Hand structures of adult males (left) and female (right) individuals photographed immediately after being in amplexus. Photographs show the dorsal side of hands (top), the paler side (middle) and a close-up of nuptial pads in males and flanged fingers in females. Scale bar represents 2mm.

### Spawn structure and development

A total of 641 *L. fletcheri* spawn were recorded over the four breeding seasons. All spawn were found in ephemeral pools, either located near the inside edge and attached to surrounding vegetation or substrate, or in the centre and free-floating. A majority of spawn were found within 24 hours of deposition (66 %), with embryos below Gosner stage 10-12. However, spawn were also found at varying times post deposition, including after 3 or 4 days when embryos were close to hatching (Gosner 20-21). Spawn less than one day old were glossy, white and horizontally oblong, possessing a semi-hemispherical structure (flat bottom and curved top surface), with a large portion of material floating up to 10-20 mm above the water’s surface. Spawn were composed of mucous secretions and a large number of medium-sized bubbles throughout (Fig. 3), which had a mean diameter of 2.2 mm (SD = 1.0, n = 42). However, the base of each spawn was comprised of a band of un-frothed mucous. Eggs were pigmented, with a mean diameter of 1.8 mm (SD = 0.2, n = 7), and were distributed irregularly throughout the spawn body, including on top of one another and near the top surface. However, no eggs were located at the base of the spawn in sections of un-frothed mucous. The mean circular diameter of fresh spawn within 24 hours of deposition was 124 mm (SD = 36, n = 468), while the mean number of eggs per spawn being 318 (SD = 206, n = 64).

**Figure 3.**
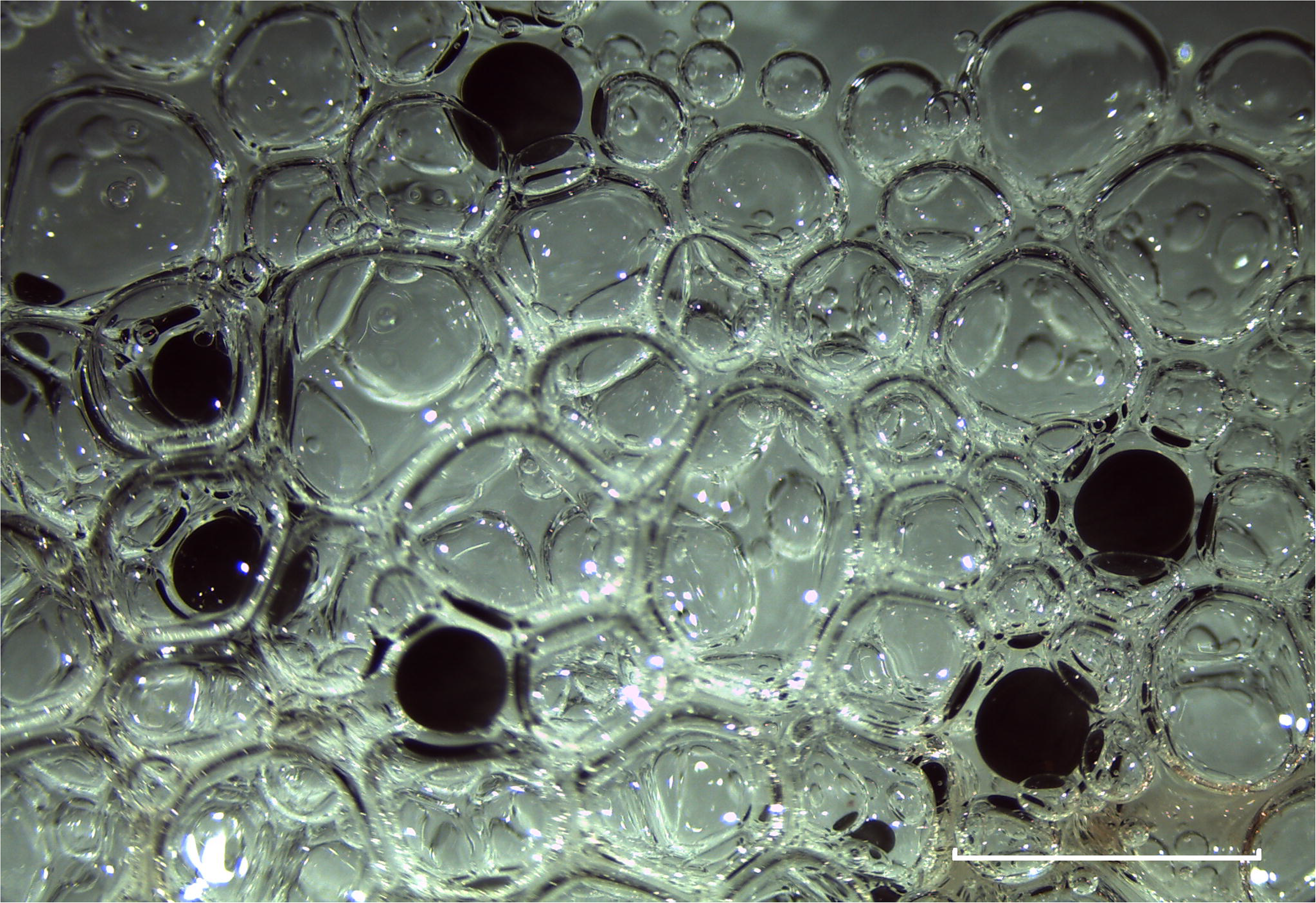
Section of a frothed spawn body made by *Lechriodus fletcheri* showing early-stage embryos interspersed through beaten mucous secretions. Spawn was imaged within a day of being oviposited. Scale bar represents 5 mm.

Over subsequent days after deposition, spawn began to flatten out and expand over the surface of the water, with the number of trapped air bubbles amongst the mucous secretions gradually declining. After 3-4 days when embryos were near hatching, spawn no longer possessed a semi-hemisphere structure but had become flat, film-like sheets of translucent mucous at the water’s surface. Few, if any, bubbles remained at this stage, with embryos distributed as a single layer at the surface. Despite this change, spawn remained floating throughout embryo development and retained some structural integrity, with a clear boundary edge and embryos remaining anchored within the surrounding mucous and not free-floating (Fig. 4). It must be stated that spawning irregularities were recorded in the field, with one spawn possessing a highly elongated, rope-like structure, and another possessing a dense collection of eggs without any surrounding frothed mucous present (Fig. 4).

**Figure 4.**
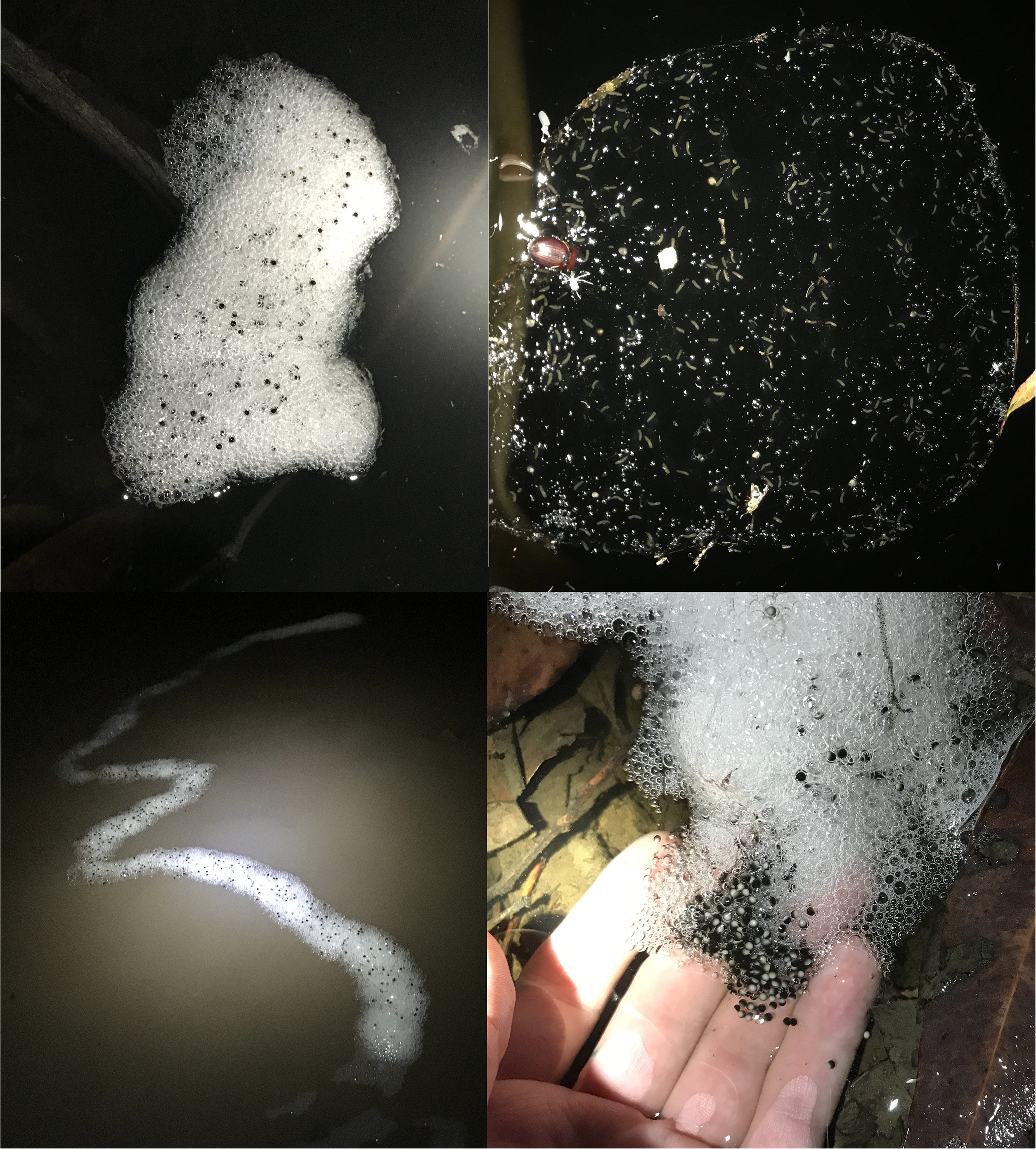
Structural and temporal differences in the frothed spawn of *Lechriodus fletcheri*. Photographs show a typical spawn within a day of being deposited (left), and after 3-4 days (top right), as well as atypical spawn structures (bottom left and right). Photographs are not to scale.

All spawn recorded in the field and laboratory showcased this gradual breakdown except for one irregular spawn which contained no residing embryos. Although similar in appearance to other *L. fletcheri* spawn, when brought back to the laboratory this egg-less spawn maintained its glossy, white appearance, semi-hemisphere structure and did not lose trapped bubbles overtime. This lack of breakdown was particularly evident when compared to a similarly aged spawn that did contain eggs, which was brought in from the field at the same time, which broke down nearly entirely after 3-4 days. The only perceived change in the egg-less spawn body was a slight reduction in overall volume as a result of desiccation (Fig. 5).

**Figure 5.**
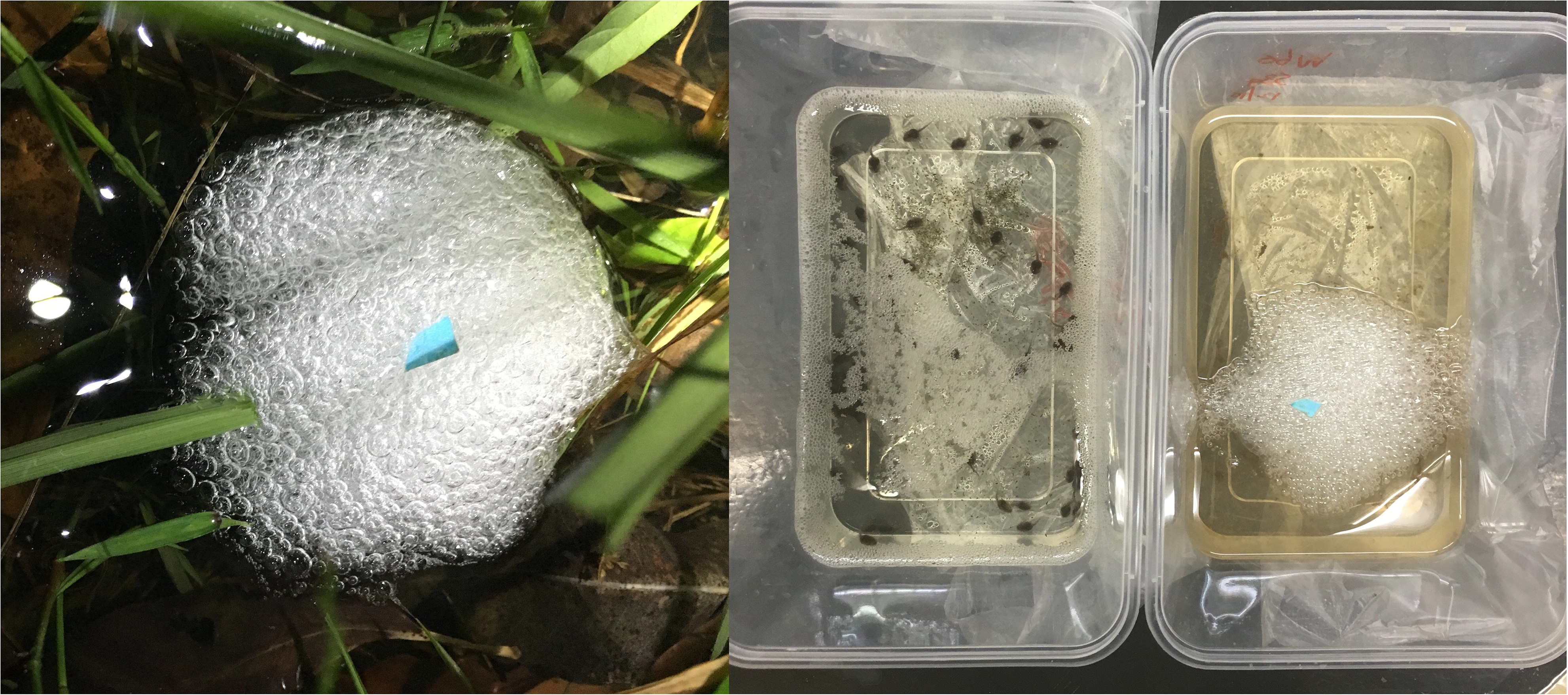
*Lechriodus fletcheri* spawn body with no residing embryos present. The left photograph shows the egg-free spawn within a day of being deposited at a breeding pool within the Watagan Mountain Range, NSW, Australia. The right photograph shows the same egg-less spawn 3-4 days after collection, alongside the remaining spawn material of a spawn body that contained embryos which was collected at the same time. Photographs are not to scale.

## Discussion

A critical aspect of ovipositional mode is the manner in which adults make a physical connection while in amplexus. In particular, the positioning of *L. fletcheri* pairs in inguinal amplexus, slightly tilted upwards in the water column, along with the location at which males grasp the females just above her hind legs, both likely aid in egg fertilization. This is because they result in the male’s cloaca sitting above and behind that of the female’s, causing the path of the eggs upon release to come close-by to the male as they rise to the surface of the water amongst the frothed mucous secretions. This connection between adult pairs is maintained through the male’s strong grip, along with nuptial pads located on the dorsal side of the two inner fingers of both hands. These pads are composed of many small, conical elevations that form a rough surface that presumably improves the male’s hold onto the female’s belly, along with a raised nub located before the inner finger on both front hands which also possess these pads.

Although *L. fletcheri* exploits shallow, ephemeral pools for reproduction, it appears that shallow water and ground contact are not required for spawn production, highlighting the possibility that stabilisation manoeuvres used by adults to maintain their position at the water’s surface may have been adapted for the purposes of froth production (Altig and McDiarmid 2007). In fact, spawn were often found free-floating near the centre of pools suggesting that the presence of structures to anchor spawn are not important for oviposition site selection. Only the female was found to facilitate the frothing of spawn, which is in agreeance with previous records by Tyler (1989). To do so, the female will complete a regular sequence of motions, initially moving her front legs back and forth out in front to agitate the water’s surface, before paddling air bubbles underneath her belly to transport them to the underside of the growing spawn. Air bubbles are captured with the assistance of fleshy flanges present on the outer two fingers of both front hands; a modification seen among Australian foam nesters (Tyler and Davies 1979). This process produces a consistent bubble spawn body similar to that created by other *Myobatrachids* of Australia, as well as *Leptodactylus* species in South America (Rivero 1969). It is a relatively simple method of froth production compared to other bubble nesting species, such as *Chiasmocleis leucost*. In that species, amplectant pairs showcase high levels of synchronised behaviour and a complex foaming sequence, diving below the water to froth the spawn from beneath by releasing air bubbles through their nostrils (Hödl and Haddad 1997). Remaining at the water’s surface may be a relatively safer method of froth spawn production that allows for rapid detection of oncoming threats in *L. fletcheri*, particularly given that amplectant pairs are already in a vulnerable state while coupled (Magnhagen 1991) and given that the splashing noises created by females while paddling may attract predators.

Fresh spawn consisted of a large number of medium to large air bubbles, approximately the same size as the neighbouring embryos, resulting in a domed-shaped spawn structure. This is in contrast to the bubble spawn produced by other species, such as *C. leucost*, which is composed of very few bubbles to form a flat film (Hödl and Haddad 1997). The number of bubbles incorporated into mucous and their size are likely to have a significant effect on the physical properties of the spawn body and, as such, its adaptive benefits for the residing embryos. For example, incorporating many large bubbles into the spawn will provide embryos with a much greater supply of oxygen that will last for a longer period without the need for diffusion from the environment, while also reducing oxygen competition between embryos that will sit further apart (Seymour and Roberts 1991; Seymour 1994; Seymour and Bradford 1995; Hödl and Haddad 1997). It will also allow a much larger spawn structure to form, particularly when compared to spawn prior to frothing, given that air bubbles in some species (e.g. *Chiromantis xerampelina*) can represent almost 80% of the spawn itself (Seymour and Loveridge 1994). This size may allow the structure to be more effective in buffering against external stresses, including sub-optimal temperatures, desiccation, and predation (Heyer 1969; Gorzula 1977; Dobkin and Gettinger 1985; Hödl 1986; Downie 1988), given its smaller surface area relative to volume that will distance eggs from the external environment (Ryan 1985; Einum *et al.* 2002). It will also cause the spawn to sit higher above the water’s surface, which may act as a physical barrier that will offer greater protection from aquatic predators. Evidence has been presented to suggest that these adaptive benefits of froth spawn production occur in *L. fletcheri*, particularly in terms of protecting eggs from cannibalism (Gould *et al.* 2019) and desiccation (J. Gould, unpublished data).

While a majority of the *L. fletcheri* spawn body is aerated, the base is composed of un-frothed secretions. Unlike species that produce foam spawn in which secretions are physically whipped (Heyer and Rand 1977; Bastos *et al.* 2010), *L fletcheri* females transport air bubbles into the growing spawn from underneath to produce bubble spawn. This lack of disturbance to the spawn body itself may cause air bubbles to gradually rise to the top of the structure, causing the base to remain un-frothed, although further investigation of this is required. Interestingly, while eggs are distributed throughout the aerated spawn, there are no eggs present within un-frothed sections. Differences in the localisation of eggs between spawn sections have been recorded in the túngara frog *(Engystomops pustulosus)*, in which a foam platform is produced prior to eggs being deposited in the centre to form a true ‘nest’ (Dalgetty and Kennedy 2010). As eggs and spawn are excreted concomitantly in *L. fletcheri*, the lack of eggs at the base is more likely to be an artefact of the process of bubble incorporation and movement to the surface. Even so, this distribution may further protect eggs from predation, given that predators will need to consume or travel through the base in order to reach the eggs above.

The *L. fletcheri* spawn body undergoes a dramatic transformation over the course of embryo development, gradually loosing incorporated air bubbles and flattening out over the surface of the water to form a translucent, thin film. This disintegration is observed across frothed nesting species but at differing rates, from less than a day (e.g. *Limnodynastes ornatus;* Tyler and Crook 1983) to several days as seen in *L. fletcheri.* Given the resultant increase in spawn surface area, this change will inevitably reduce the ability of the spawn body to buffer eggs from external stresses. This appears counterintuitive at first glance as both early- and late stage embryos are equally as vulnerable to external stresses, which should select for the structure to remain stable throughout development. However, this change may be adaptive, as an increase in spawn surface area will result in each embryo being in direct contact with the surrounding water, which may compensate for increases in oxygen demand by later stage embryos (Seymour and Roberts 1991) or prevent embryos from becoming trapped within the spawn upon hatching. Not only does this suggest the temporal shift in spawn structure is deliberate and driven by changes in the needs of the embryos over the course development, but indicates a trade-off between the ability of the spawn to protect embryos from environmental stresses and to prevent anoxia or allow for successful hatching.

Interestingly, one spawn body collected from the field without eggs did not follow this transition into film, remaining stable in appearance for days. Similar observations have been recorded in the frothed spawn of *E. pustulosus* in the absence of developing embryos (Fleming *et al.* 2009). In *L. fletcheri*, this could indicate that the frothed fluid that makes up the spawn body is relatively stable and that its gradual breakdown is not random but a controlled process orchestrated by the developing embryos themselves. Embryos of other amphibians species are known to secrete active substances into the perivitelline fluid to facilitate their hatching from the egg (Salthe 1965; Urch and Hedrick 1981), with those of foam nesting *Chiromantis* are known to release enzymes that degrade surrounding foam to allow for their release from the spawn upon hatching (Coe 1974). As such, I hypothesise that *L. fletcheri* embryos also release substances that contribute to the breakdown of the spawn body overtime, as a deliberate adaptation to modify their immediate environment to ensure conditions remain suitable for their changing needs.

It is clear that the frothed spawn body is a dynamic medium for embryo development in *L. fletcheri*, with temporal changes in its structure facilitating the changing needs of the embryos up until hatching. While ovipositional mode is singular in this species, irregular spawn structures are present within the population, including the production of elongated spawn that form strings and egg clutches lacking in surrounding frothed mucous secretions. However, these irregularities are more likely to be attributed to disturbances of amplectant pairs at the egg deposition site rather than representations of alternative oviposition modes. Given that froth spawn production and breakdown differ immensely between species, and the adaptive benefits these structure provide to embryos, precise ovipositional data needs to be collected for each species to assist in determining how they have evolved to become reproductively suited to their respective environments.

## Acknowledgements

This work was conducted under NPWS Scientific license (no. SL101991) and approved by the University of Newcastle Animal Care and Ethics Committee (no. A-2011-138). All experimental procedures were performed in accordance with the Australian code for the care and use of animals for scientific purposes. This research did not receive any specific funding.

